# Microbiome divergence of marine gastropod species separated by the Isthmus of Panama

**DOI:** 10.1101/2021.07.08.451645

**Authors:** Alexander T. Neu, Mark E. Torchin, Eric E. Allen, Kaustuv Roy

**Affiliations:** Section of Ecology, Behavior and Evolution, Division of Biological Sciences, University of California San Diego, La Jolla, CA, USA; Smithsonian Tropical Research Institute, Balboa, Ancon, Republic of Panama; Section of Molecular Biology, Division of Biological Sciences, University of California San Diego, La Jolla, CA, USA; Marine Biology Research Division, Scripps Institution of Oceanography, University of California San Diego, La Jolla, CA, USA

**Keywords:** host-microbiome, Isthmus of Panama, geminate species, divergence, vicariance

## Abstract

The rise of the Isthmus of Panama ~3.5 mya separated populations of many marine organisms, which then diverged into new geminate sister species currently living in the Eastern Pacific Ocean and Caribbean Sea. However, we know very little about how such evolutionary divergences of host species have shaped their microbiomes. Here, we compared the microbiomes of whole-body and shell-surface samples of geminate species of marine gastropods in the genera *Cerithium* and *Cerithideopsis* to those of congeneric outgroups. Our results show that the effects of the Isthmus on microbiome composition varied among host genera and between sample types within the same hosts. In the whole-body samples, microbiome compositions of geminate species pairs in the focal genera tended to be similar, likely due to host filtering, although the strength of this relationship varied among the two groups and across similarity metrics. Shell-surface communities showed contrasting patterns, with co-divergence between the host taxa and a small number of microbial clades evident in *Cerithideopsis*, but not *Cerithium*. These results suggest that (i) the rise of the Isthmus of Panama affected microbiomes of geminate hosts in a complex and clade-specific manner and (ii) host-associated microbial taxa respond differently to vicariance events than the hosts themselves.

## Introduction

The formation of the Isthmus of Panama (IP) ~3.5 mya [1,2] led to a mixing of terrestrial faunas between North and South America, commonly known as the Great American Biotic Interchange [3,4]. In the marine environment, however, the rise of the Isthmus separated populations of many species, which then evolved to form geminate species pairs [5], with one currently inhabiting the nutrient-rich upwelling zone of the Eastern Pacific Ocean and the other inhabiting the warmer, more oligotrophic Caribbean Sea [6]. Geminate species pairs provide a powerful set of natural evolutionary experiments that have been used to investigate how closely related species evolve under different environmental conditions [7–9] and to better constrain rates of molecular evolution [e.g., 1,10,11]. However, it remains unclear how the microbiomes of geminate hosts have diverged in response to this vicariance event. Although there is a framework for examining how biogeographic isolation and environmental change may structure the microbiomes of geminate hosts [12], host-associated microbial divergence across the IP has so far only been studied using eggs of geminate species of echinoids [13].

Tests of concordance between divergence patterns of microbiomes and those of their hosts rely on correlations between the phylogenetic relatedness of host taxa and the similarity of their microbiomes, a pattern termed “phylosymbiosis” [14,15]. Under this model, geminate taxa are expected to harbor microbial communities that are significantly more similar to each other than to more distantly related hosts currently living in the same environment. This pattern may arise either as the result of co-phylogenetic divergence between hosts and their associated microbes, host filtering, or both [15]. In the first case, geminate hosts and their microbiomes (or a subset thereof) are expected to diverge in concert with one another after the closure of the IP, resulting in co-phylogenetic relationships between the geminate host clades and associated microbial taxa. In the second case, aspects of the host environment such as morphology, immune function and diet (e.g., nutrient and metabolite availability) may exclude certain microbial taxa while providing highly suitable conditions for others [16]. Since closely related hosts, such as geminates, often share many traits, they would be predicted to select for more compositionally similar microbial communities. While they are distinct processes, co-speciation and host filtering are not mutually exclusive pathways to phylosymbiosis, and have been shown to act in concert to generate concordance between host evolutionary history and microbiome composition in some host groups, such as corals [17].

Here we use geminate species from two genera of marine gastropods, *Cerithium* and *Cerithideopsis,* along with congeneric outgroups in each case, to test for phylosymbiosis resulting from vicariance due to the rise of the IP. Among our focal host species, *Cerithium lutosum* and *Cerithium stercusmuscarum* form a well-supported geminate species pair, with *Cerithium atratum* as an outgroup [11; Fig. 1]. *Cerithideopsis mazatlanica* and *Cerithideopsis valida* diverged in the Eastern Pacific after the rise of the IP and are a geminate clade to *Cerithideopsis pliculosa* in the Caribbean, with the Eastern Pacific *Cerithideopsis montagnei* as a congeneric outgroup [11; Fig. 1]. Both outgroups were sampled in the same geographic locations as our geminate taxa (Fig. 1). The *Cerithium* species used here are typically found in near-shore environments with close proximity to mangroves. *Cerithium stercusmuscarum* largely occupies rocky shores in the Eastern Pacific, while its geminate *Cerithium lutosum* and the outgroup *Cerithium atratum* are found in coral rubble or sandy flats in the Caribbean. The *Cerithideopsis* species sampled here all occupy shallow, muddy environments and associated mangrove habitats [11]. Though all of these species forage in the mud, *Cerithideopsis montagnei* differs from its congeners by climbing mangrove roots during high tides.

**Figure 1.**
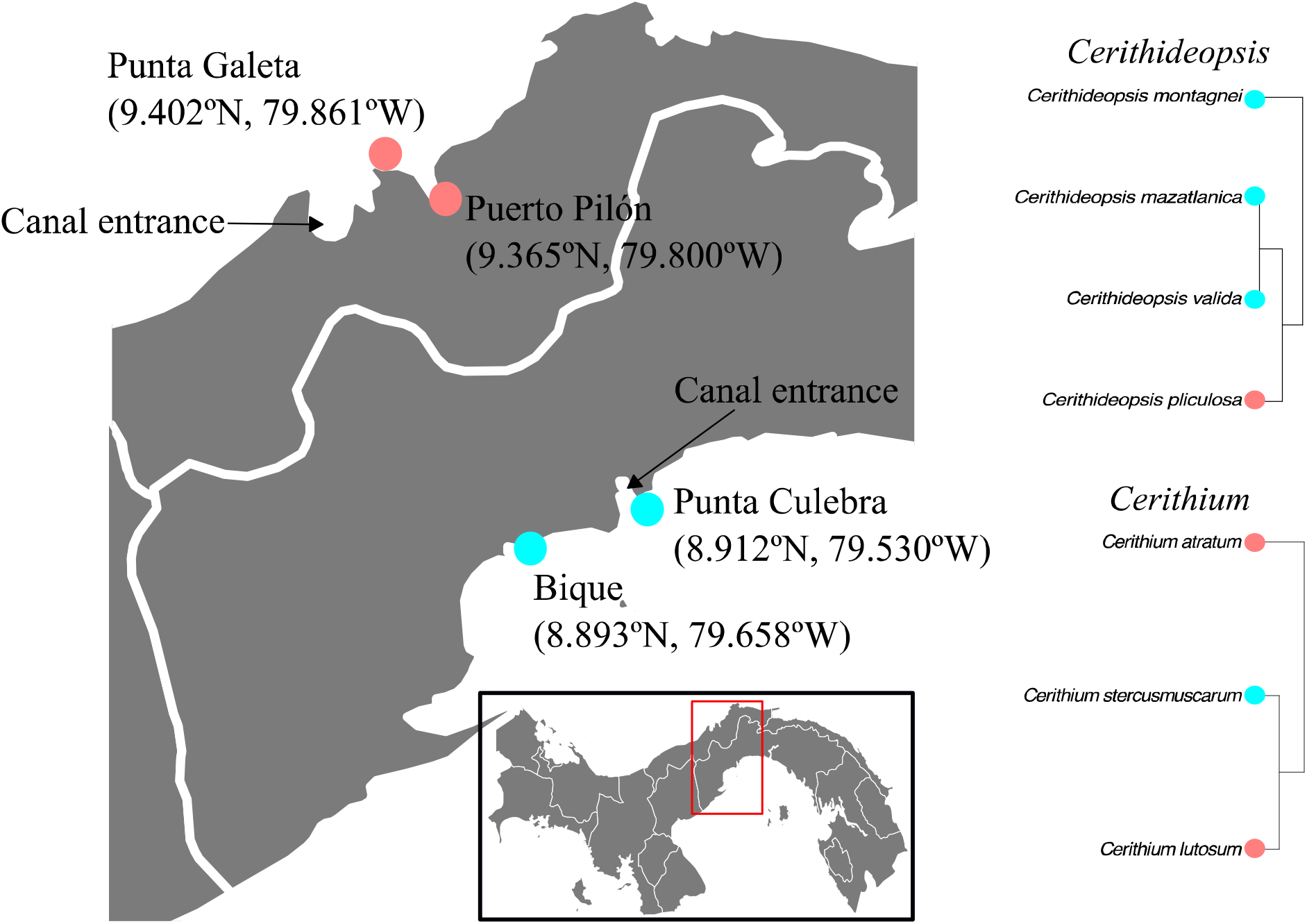
Map of sampling area. The map of Panama (bottom inset) shows the location of the detailed map of the sampling area (left). Blue dots indicate sampling sites in the Eastern Pacific, while pink dots indicate sampling sites in the Caribbean. Phylogenies of the species sampled from each host genus (right) have colors to represent the side of the IP each species was sampled from.

For each host, we used soft body tissues (hereafter “whole-body”) as well as swabs of the shell-surfaces of multiple individuals (Table S1) to test the hypothesis that geminate species pairs would harbor microbiomes that are both compositionally and phylogenetically more similar to one another than to their sympatric congeners.

## Methods

### Sampling

We sampled multiple individuals of each of seven different marine gastropod species from our focal genera at four sites across the IP (Figure 1, Table S1). We swabbed the shell-surface of each individual in the field using a sterile 152mm swab (Grenier Bio-One, Frickenhausen, Germany), avoiding any visible patches of mud. At each site, we also sampled the top 3 cm of sediment in a sterile 50 mL conical tube and collected two replicate seawater samples by filtering 500mL of surface seawater through 0.22μm Sterivex filters (MilliporeSigma, Burlington, MA, USA). Samples were transported on ice to the Naos Marine Laboratory at the Smithsonian Tropical Research Institute (STRI), where they were kept at − 20°C. Subsequently, the whole-body of each individual was removed from the shell, triple-rinsed with 95% EtOH and preserved at −20°C. Samples were later shipped to the University of California, San Diego for DNA extraction and sequencing.

### Sequencing

We used the Qiagen Blood and Tissue kit (Qiagen GmbH, Hilden, Germany) to extract DNA from our samples, following the manufacturer’s instructions. We used the protocol of [18] to amplify and purify the V4 region of the 16S rRNA gene. The resulting amplicons were sequenced on the MiSeq platform (Illumina, 2×250bp output) at the DNA Technologies Core, University of California Davis. In order to verify host identities, we also amplified the cytochrome oxidase *c* subunit I (COI) gene from one individual of each host species using the PCR protocol of [11]. The PCR products were cleaned using an Exo-SAP approach and Sanger sequenced at Eton Bioscience, Inc. (San Diego, CA, USA).

### Data Analyses

We used ‘DADA2’ v1.18.0 [19] in R v4.0.3 [20] to process the 16S rRNA gene sequences. The resulting amplicon sequence variants (ASVs) were assigned taxonomy using the Silva database v138 [21] and all sequences not matching Bacteria or Archaea, as well as any chloroplasts and mitochondria, were removed. We aligned the remaining sequences using MAFFT v7.310 [22] and constructed microbial phylogenies for further analyses with FastTree v2.1.11 [23] using the GTR + CAT model and midpoint rooting.

For each host genus, we tested for differences in alpha diversity among the hosts using two sets of metrics. First, we used the entire, unrarefied microbial dataset to estimate Hill numbers with q of 0 and 1 for each sample using the package hilldiv v1.5.1 [24]. Using Hill numbers with q = 0 does not account for relative abundance and is equivalent to the number of observed ASVs, while q = 1 weights each ASV by its relative abundance and is equivalent to the exponential of Shannon’s index (H’) [25]. Second, we generated 100 ASV tables, each rarefied to 3,000 sequences per sample, using the package metagMisc v0.0.4 [26]. We then calculated the number of observed ASVs and Shannon’s H’ for each sample from each ASV table and used the average values to account for any differences in individual random draws. In both cases, we tested for differences between host species using Kruskal-Wallis tests and further investigated significant differences using Dunn’s tests in the package dunn.test v1.3.5 [27].

All subsequent analyses used both the unrarefied dataset, normalized via total sum scaling (TSS) [30], as well as the 100 rarefied datasets (3,000 sequences/sample), unless otherwise specified. We tested for differences in microbiome composition between host species using permutational multivariate analyses of variance (PERMANOVAs) in the package phyloseq v1.34.0 [29] with Bray-Curtis Dissimilarity (BCD) and unweighted UniFrac Distance [30] as our beta diversity metrics. We use both of these metrics in our analyses as they emphasize different attributes; BCD includes relative abundance information, while unweighted UniFrac includes a measure of phylogenetic relatedness of the sampled microbial taxa. We also conducted PERMANOVAs on the TSS dataset to determine whether the host-associated microbiomes were significantly different from the microbial communities in the sediment and water samples. We tested for homogeneity of dispersions between species and sample types using the betadisper function in vegan v2.5-7 [31] for all iterations.

To investigate the role of host phylogenetic distance on microbiome composition more directly, we pooled together all microbial sequences from each individual host species. Since *Cerithium stercusmuscarum* was collected from two sites, we pooled all sequences from this species together but also pooled each site individually. We then clustered the samples using unweighted paired group with arithmetic means (UPGMA), using both BCD and unweighted UniFrac, and the TSS and rarefied datasets. To test the robustness of clustering, we calculated Dunn’s Index for each node using the package clValid v0.7 [32]. We used the normalized Robinson-Foulds metric (nRF) in the package phangorn v2.5.5 [33] to test for congruency between the UPGMA plots and the host phylogeny generated from COI sequences using FastTree v2.1.11 [23] with the GTR + γ model. For some samples, relationships varied across different rarefied datasets, so we generated boxplots showing the BCD and unweighted UniFrac distance between species across all 100 rarefied datasets.

To identify microbial clades contributing to similarities between geminate host species, we generated network plots using sparse partial least squares discriminant analysis (SPLS-DA) in the package mixOmics v6.14.0 [34] at taxonomic levels ranging from ASV to class. This analysis requires a log-ratio transformation and was only conducted on the unrarefied dataset using a centered log-ratio transformation. Microbial taxa that were positively correlated with the geminate species (r > 0.6) or negatively correlated with the congeneric outgroups (r < −0.6) were considered to contribute to phylosymbiosis resulting from the rise of the IP [35]. To further investigate whether the taxa contributing to the differentiation between the geminate clades and outgroups also demonstrated potential co-phylogenetic relationships with the hosts, we used the parafit function in ape v5.4.1 [36].

## Results

### Alpha Diversity

#### Cerithium

The whole-body microbiomes of all three species of *Cerithium* showed similar Hill numbers at q = 0 (Fig. 2a) and observed ASVs after rarefying (Fig. S1a), and there were no significant differences in total ASV richness between the geminate and outgroup species (Tables S2, S3). However, at q = 1, the Hill numbers of *C. atratum* were significantly higher than those of the geminate pair *C. lutosum* and *C. stercusmuscarum* (Fig. 2c, Table S2), as confirmed by Shannon’s H’ values in the rarefied dataset (Fig. S2a, Table S3). In the *Cerithium* shell-surface microbiome, no significant differences were found between geminate and outgroup species using any of the alpha diversity metrics (Figs. 2b, 2d, S1b, S2b, Tables S2, S3).

**Figure 2.**
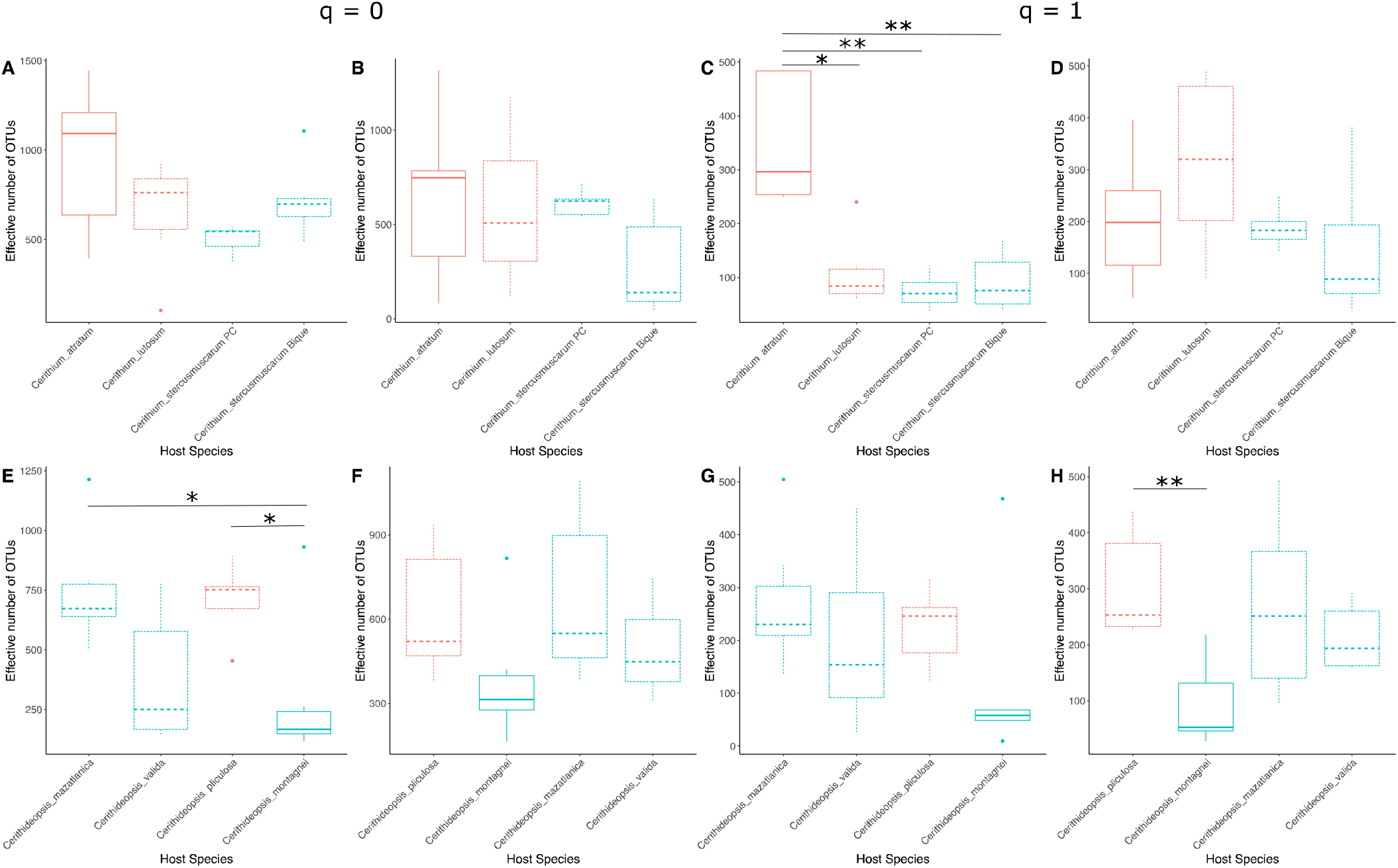
Hill numbers of q = 0 (A,B,E,F) and q = 1 (C,D,G,H) from whole-body (A,C,E,G) and shell-surface (B,D,F,H) samples. Blue boxes represent samples collected from the Eastern Pacific and pink boxes represent samples collected from the Caribbean. Dashed boxes represent geminate species and solid boxes represent congeneric outgroups. Asterisks indicate samples which were found to be significantly different by Dunn’s test at p < 0.05 (*) or p < 0.005 (**).

#### Cerithideopsis

In this group, whole-body samples showed significant differences in Hill numbers at q = 0 (Fig. 2e; Table S2) and marginally significant differences in observed ASV richness (Fig. S1c, Table S3). This was driven by the low diversity of *C. montagnei*, which was significantly different from *C. mazatlanica* and *C. pliculosa* (Dunn’s tests: *p < 0.05*), but not *C. valida*. Interestingly, the geminate and outgroup species did not show significant differentiation at q = 1 (Fig. 2f; Table S2) or using Shannon’s index (Fig. S2c, Table S3), largely due to the presence of a single *C. montagnei* sample with high diversity (Fig. 2e, Supplement 2). *C. montagnei* showed significantly lower diversity than *C. mazatlanica* and *C. pliculosa* (Dunn’s test: *p = 0.0084* and *p = 0.005*, respectively), but not *C. valida*, when this high diversity individual was removed. In the *Cerithideopsis* shell-surface swabs, Hill numbers at q = 0 did not show significant differences between the host species (Fig. 2g; Table S2), again due to a single high diversity sample of *C. montagnei* (Fig. 2g, Supplement 2), though the rarefied observed ASV count showed marginally significant differentiation (Fig. S1d, Table S3). Removing the high diversity sample from the Hill number analyses resulted in significant differences between *C. montagnei* and the higher diversity *C. pliculosa* and *C. mazatlanica* (Dunn’s test: *p = 0.0129* and *p = 0.0135*, respectively), but not *C. valida*. When q = 1, significant differentiation (Table S2) was driven by the difference between *C. pliculosa* and *C. montagnei* (Dunn’s test: *p = 0.0244*), as *C. mazatlanica* and *C. valida* were not significantly different from either of these hosts (Fig. 2h). These results were confirmed by Shannon’s H’ from the rarefied dataset (Fig. S2d, Table S3).

### Taxonomic Composition

#### Cerithium

Whole-body samples of individual host species were largely comprised of Actinobacteriota, Alphaproteobacteria, Cyanobacteria and Gammaproteobacteria (Fig. 3a). Interestingly, the most common group found in the geminate species *C. lutosum* and *C. stercusmuscarum* was “Unclassified Bacteria”, as determined by the Silva database. A single ASV within this group (ASV10677) comprised, on average, greater than 20% of the relative abundance of the microbiome in these hosts. The closest named relative to this microbe is a member of the phylum Chlorobi, *Chloroherpeton thalassium* (81.89% sequence similarity using NCBI BLASTn). The two populations of *C. stercusmuscarum* whole-body samples were differentiated by the relative abundances of the phyla Crenarchaeota, which was higher at Bique, and Cyanobacteria, which was higher at Punta Culebra (Fig. 3a). *Cerithium* shell-surface microbiomes were comprised mainly of Alphaproteobacteria, Bacteroidota, Cyanobacteria, Gammaproteobacteria and Planctomycetota (Fig. 3b).

**Figure 3.**
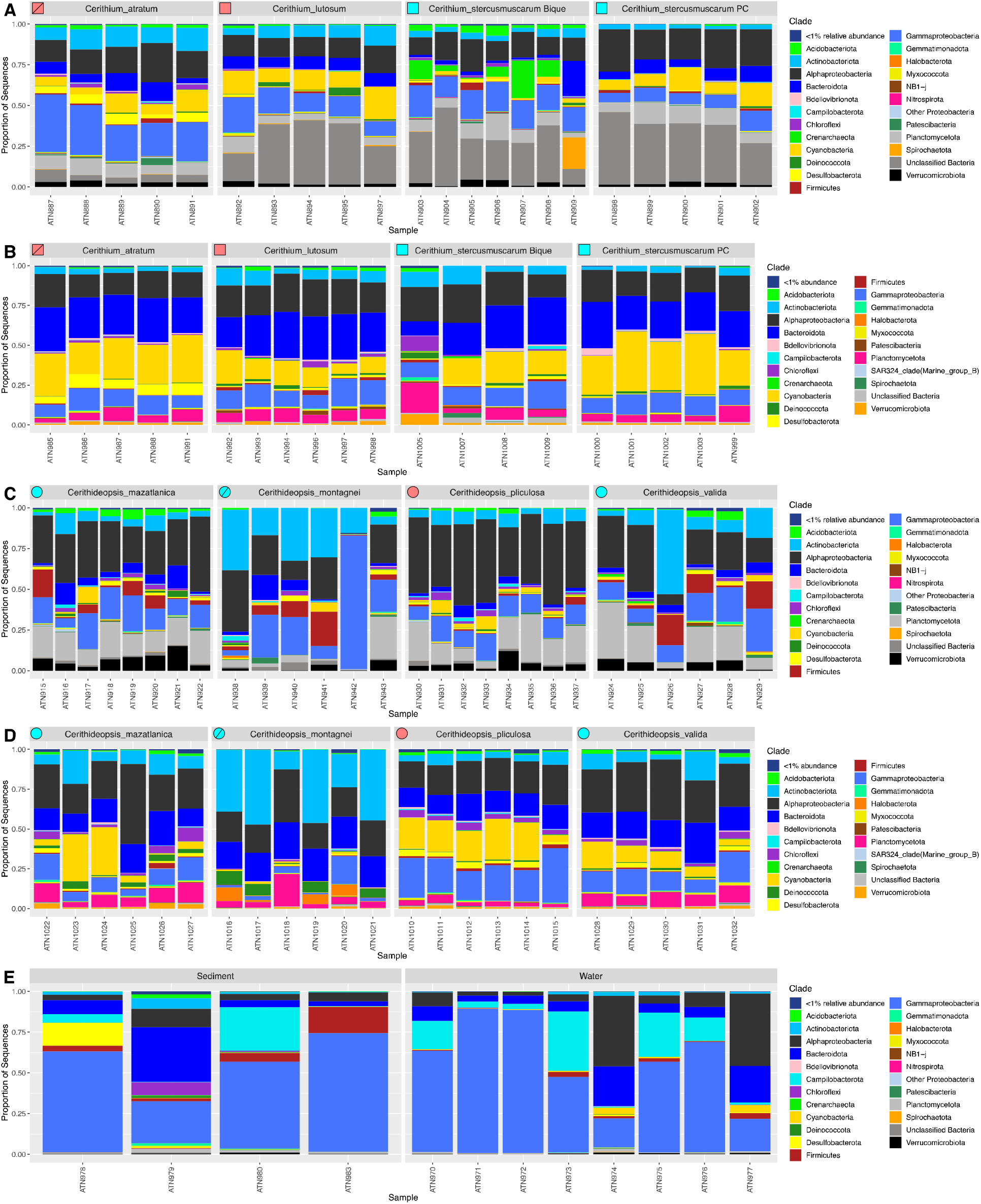
Barplots of the relative abundances of microbial phyla in the whole-body (A,C), shell-surface (B,D) and environmental (E) samples. The phylum Proteobacteria is separated into classes for increased resolution. In the top-left corner of each plot, the shape refers to the genus (squares and circles for *Cerithium* and *Cerithideopsis*, respectively), the color refers to the ocean basin of sampling (blue and pink for Eastern Pacific and Caribbean, respectively) and geminate species are open while congeneric outgroups are represented by a diagonal line.

#### Cerithideopsis

Whole-body samples of these host species contained high relative abundances of Alphaproteobacteria, Bacteroidota and Gammaproteobacteria. However, *C. montagnei* showed higher abundance of Actinobacteriota while the geminate clade including *C. mazatlanica*, *C. valida* and *C. pliculosa* had increased relative abundances of Planctomycetota and Verrucomicrobiota (Fig. 3c). *Cerithideopsis* shell-surface microbiomes showed high relative abundances of Alphaproteobacteria, Bacteroidota, Gammaproteobacteria and Planctomycetota (Fig. 3d). However, *C. montagnei* shell-surfaces showed higher levels of Actinobacteriota and a notable decrease in Cyanobacteria compared to the geminate species (Fig. 3d).

Microbial community compositions of sediment and water samples varied among the collection sites, but nearly all were dominated by members of the class Gammaproteobacteria and were compositionally distinct from the microbiomes of all host species (Fig. 3e).

### Host evolutionary history and microbiome composition

Results of PERMANOVAs show that in both host genera, the microbiome compositions of individual host species are distinct, regardless of tissue type, normalization method or beta diversity metric used (Tables S4, S5), and significantly different from the environmental samples (Table S6).

#### Cerithium

UPGMA clustering of whole-body samples was consistent with the phylosymbiosis hypothesis when using BCD and either normalization method, with microbiomes of geminate species *C. lutosum* and *C. stercusmuscarum* being compositionally more similar to each other than to *C. atratum* (Figs. 4a, S3a). However, when *C. stercusmuscarum* individuals were separated by collection site, *C. stercusmuscarum* samples from Punta Culebra were more similar to *C. lutosum* than to the *C. stercusmuscarum* samples from Bique (Fig 5a). Analyses using unweighted UniFrac showed a different pattern, with *C. lutosum* microbiomes clustering closest to those from *C. atratum* with the *C. stercusmuscarum* populations as outgroups (Figs. 4a, 5a, S4a). UPGMA clustering of shell-surface samples from *Cerithium* did not match the host phylogeny, regardless of distance metric or normalization method (Figs. 4b, 5b, S3b, S4b). Instead, microbiome composition appeared to be determined by the local environment, as the Caribbean species *C. lutosum* and *C. atratum* were more similar to each other than to either population of the Eastern Pacific *C. stercusmuscarum*.

**Figure 4.**
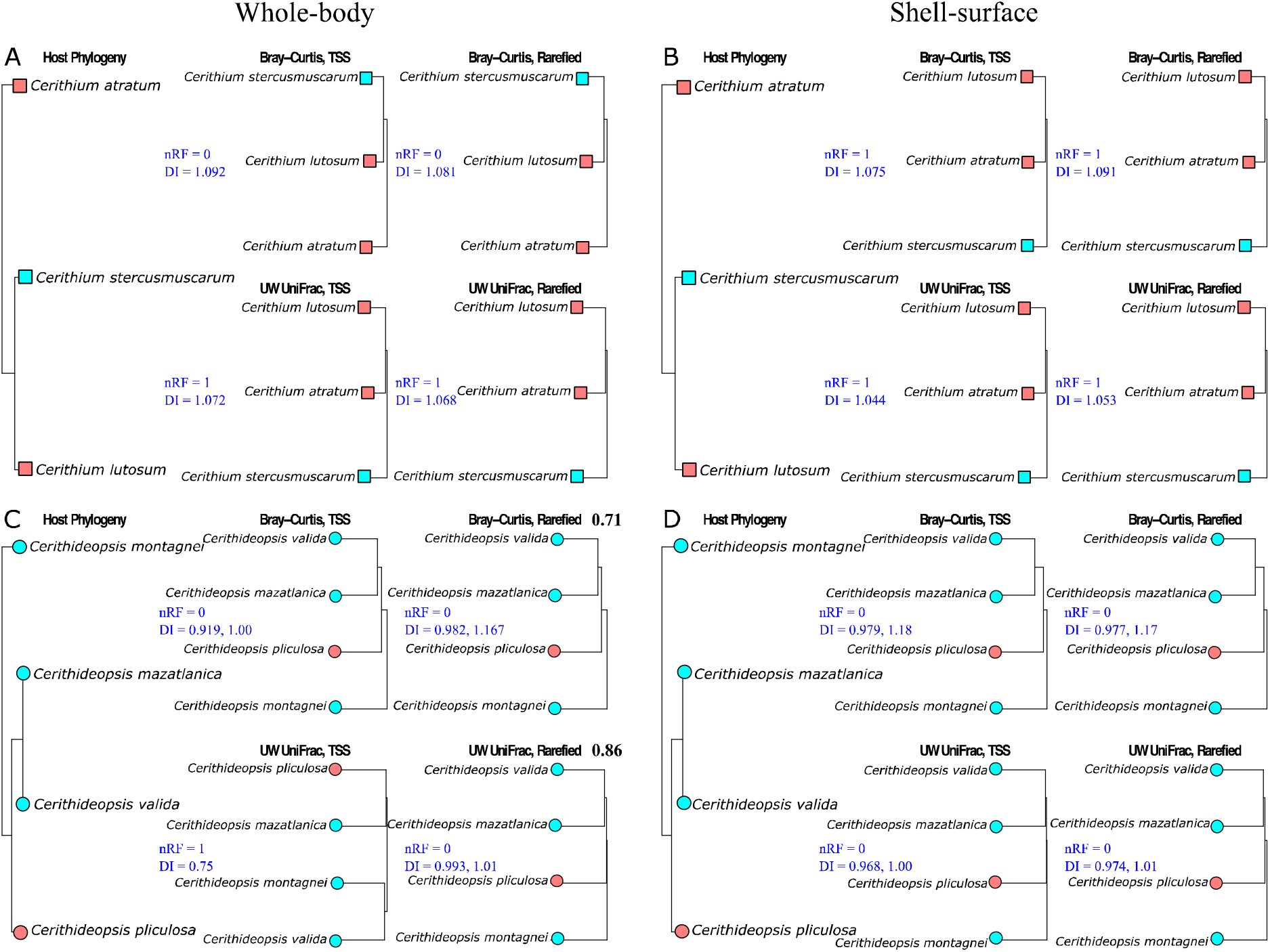
Comparing host phylogeny to microbiome composition in the whole-body (A,C) and shell-surface (B,D) communities of *Cerithium* (A,B) and *Cerithideopsis* (C,D) species. For each sample type, we show the host phylogeny (left) and UPGMA plots generated using BCD and TSS (top middle), BCD and rarefying (top right), unweighted UniFrac and TSS (bottom middle) and unweighted UniFrac and rarefying (bottom right). Each UPGMA plot includes the normalized Robinson-Foulds value (nRF) and Dunn’s index (DI) in blue. Multiple DI values provide the clustering strength when the UPGMA plot is divided into two and three clusters, respectively. Bolded numbers in (C) indicate the proportion of rarefied datasets which produce the pattern shown. Tip shape refers to the genus (squares and circles for *Cerithium* and *Cerithideopsis*, respectively) and color refers to the ocean basin of sampling (blue and pink for Eastern Pacific and Caribbean, respectively).

**Figure 5.**
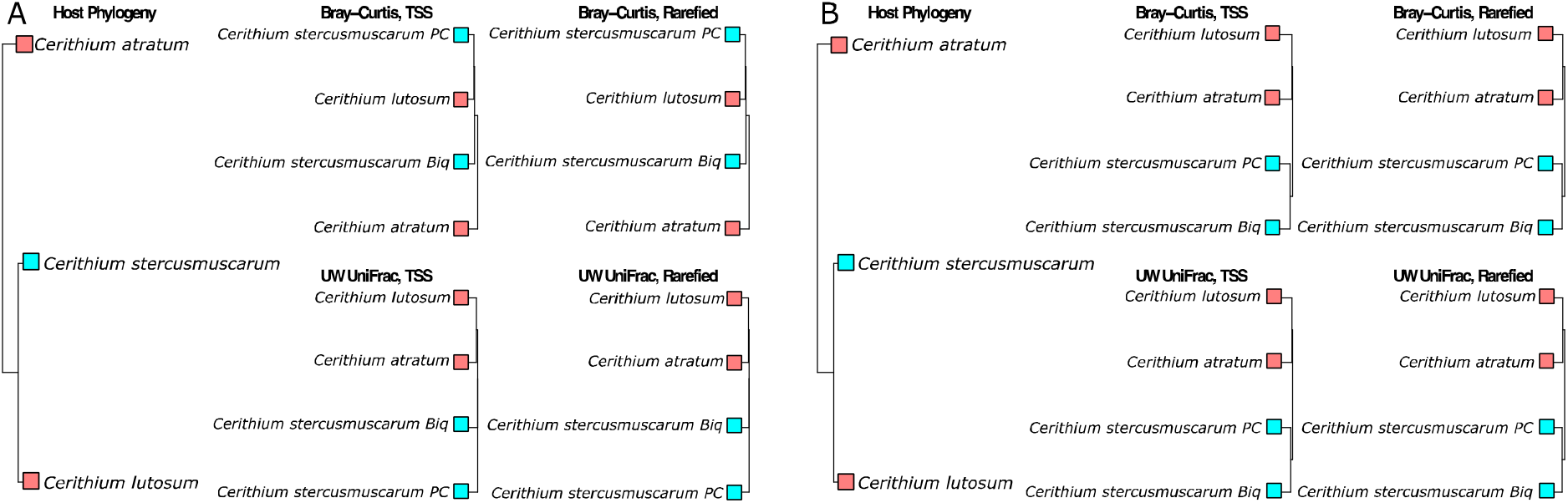
Comparing host phylogeny to microbiome composition in the whole-body (A) and shell-surface (B) communities of *Cerithium* species, separated by collection site. For each sample type, we show the host phylogeny (left) and UPGMA plots generated using BCD and TSS (top middle), BCD and rarefying (top right), unweighted UniFrac and TSS (bottom middle) and unweighted UniFrac and rarefying (bottom right). Tip color refers to the ocean basin of sampling (pink and blue for Eastern Pacific and Caribbean, respectively).

#### Cerithideopsis

When using BCD and the TSS dataset, UPGMA clustering of host microbiome compositions mirrored the host phylogeny (Fig. 4c). However, BCD of samples rarefied to 3,000 sequences produced mixed results depending on which of the 100 resampled ASV tables was used (Fig. 4c). In 71% of the datasets, results were consistent with the hypothesis of phylosymbiosis, but in the other 29% of cases *C. valida* microbiomes were shown to be more similar to those of *C. montagnei* than to those of its sister species, *C. mazatlanica*, or the geminate *C. pliculosa*. This appears to be driven by the compositional dissimilarity between individuals of *C. valida*, as the BCD between this species and other *Cerithideopsis* encompass a large range of values (Fig. S3c). Clustering the TSS dataset using unweighted UniFrac distance does not support the phylosymbiosis hypothesis. However, the rarefied OTU tables once again presented split patterns, with 86% of the results being consistent with predicted phylosymbiosis (Fig. 4c, S4c). In contrast to the whole-body samples, shell-surface microbiomes of the *Cerithideopsis* geminate clade consistently clustered together, with *C. montagnei* as the outgroup, regardless of the distance metric or normalization method used (Figs. 4d, S3d, S4d).

### Unique microbial associates of geminate hosts

#### Cerithium

Only one ASV (ASV10677) significantly differentiated whole-body samples of *C. lutosum* and *C. stercusmuscarum* from *C. atratum* (Fig. S5). As discussed above, this ASV was the most dominant member of the *C. lutosum* and *C. stercusmuscarum* communities but was not highly abundant in the *C. atratum* microbiome (Fig. S5). In the shell-surface communities, no individual taxa could significantly differentiate the geminate pair from *C. atratum*.

#### Cerithideopsis

These samples contained several microbial taxa that significantly differentiated the geminate clade of *C. mazatlanica*, *C. valida* and *C. pliculosa* from the outgroup, *C. montagnei*. In the whole-body, two genera and four families were significant drivers of this differentiation (Fig. S6, Table S7), while shell-surface swabs contained four genera, five families, four orders and two classes of microbes which separated the geminates from *C. montagnei* (Fig. S7, Table S7).

### Host-microbe co-phylogenetic relationships

#### Cerithium

We were unable to test for potential co-phylogenetic relationships between *Cerithium* samples and the microbial clades driving differentiation between geminates and the outgroup, as only a single ASV was responsible for this pattern in the whole-body samples, and none differed significantly between the geminate and outgroup shell-surface samples (Table S7).

#### Cerithideopsis

Whole-body samples of this genus did not show any co-phylogenetic relationships across multiple levels of microbial taxonomy (Table S7). In the shell-surface swabs, however, two families (A4b and Thermoanaerobaculaceae), two orders (SBR1031 and Thermoanaerobaculales) and the class Gammaproteobacteria all showed significant co-phylogenetic relationships with the host (Table S7).

## Discussion

Whole-body and shell-surface samples of tropical marine gastropod hosts from the genera *Cerithium* and *Cerithideopsis* house diverse, complex and host-specific microbial communities. The richness of ASVs was surprisingly similar across host genera in both the whole-body and shell-surface samples, with the exception of relatively low diversity in *Cerithideopsis montagnei*. This may reflect the unique ecology of *C. montagnei*, which forages in the mud like the other *Cerithideopsis* species, but climbs mangroves during high tide, potentially reducing its exposure to taxa present in the seawater that surrounds its congeners, resulting in reduced diversity. Differences in microbial diversity between the geminate taxa and their congeneric outgroups are most evident using diversity metrics that account for relative abundances. The geminate pair *Cerithium lutosum* and *C. stercusmuscarum* had significantly lower Hill numbers at q = 1 and Shannon’s H’ compared to their congeneric outgroup, *C. atratum*. These metrics suggest that fewer taxa dominate the microbiomes of these geminate species as compared to *C. atratum*. Within *Cerithideopsis*, the trend is reversed, with *C. montagnei* harboring reduced diversity compared to its geminate congeners using these metrics.

Compositionally, the whole-body and shell-surface microbiomes of *Cerithium* and *Cerithideopsis* species were distinct from one another but were comprised of taxa commonly associated with marine invertebrate hosts. Many of the dominant microbial clades in the whole-body samples, including Alphaproteobacteria, Gammaproteobacteria and Planctomycetota, have been found previously in marine molluscan tissues [18,37]. The shell-surfaces of *Cerithium* and *Cerithideopsis* species also contained a number of clades previously found on shells of other marine invertebrates, including Alphaproteobacteria, Bacteroidota and Cyanobacteria [38,39]. In contrast, our environmental samples were largely dominated by Gammaproteobacteria, which are commonly found in seawater and intertidal sediment communities [40,41]. Thus the microbiome compositions of all of our host species are distinct from that of the ambient environment, as is common in marine invertebrates [e.g., 36,43].

Similarities of microbiomes of geminate species varied considerably depending on the host genus, the type of sample and the metric used. *Cerithium* whole-body samples conformed to a phylosymbiosis pattern when using BCD, as *C. lutosum* was more similar to its geminate *C. stercusmuscarum* than to *C. atratum* samples from the same environment. However, when using unweighted UniFrac, *C. lutosum* clustered more closely with *C. atratum* than *C. stercusmuscarum*. These results suggest that the relative abundances of certain microbial taxa are more similar between the geminate species than the congeneric outgroup, while microbial taxa occurring on the same side of the Isthmus tend to be closer evolutionarily. A single ASV, ASV10677, was primarily responsible for the similarity between the geminates *Cerithium lutosum* and *C. stercusmuscarum* observed in the BCD dataset. Although this ASV was present in all three *Cerithium* hosts, its relative abundance was nearly an order of magnitude higher in the geminates when compared to *C. atratum*. ASV10677 was also found in all of our sediment and water samples, but at <1% relative abundance, suggesting selective enrichment in *C. lutosum* and *C. stercusmuscarum*. The reason for this is unclear, though the closest named relative to this taxon is *Chloroherpeton thalassium*, a green sulfur bacterium [43], suggesting a potential role in sulfur cycling. The *Cerithium* geminate species are found associated with mangrove sediments, where sulfur cycling occurs [44], while *C. atratum* is often found further out on the reef flat.

When the two sampled populations of *Cerithium stercusmuscarum* were analyzed separately, the BCD dataset showed a different pattern. Though the geminate species were separated from the congeneric outgroup, *C. atratum*, *C. stercusmuscarum* populations did not cluster together, potentially due to their positions on opposite sides of the Panama Canal (Fig. 1). This shows that intraspecific, site-level differences in microbiome composition can be greater than the differences observed between species and should be accounted for in future studies.

Shell-surface microbiomes of *Cerithium* did not show any significant geminate signal, regardless of the metric used. Thus, the ambient environment may have a greater impact on shell-surface microbiome composition in this group than evolutionary history, especially as *C. atratum* and *C. lutosum* were collected from the same site.

In the whole-body samples of *Cerithideopsis*, compositional similarities of geminate hosts varied depending on the method used. In the TSS dataset, geminate species clustered together if BCD was used, but not unweighted UniFrac, a result similar to that in *Cerithium*. In our rarefied datasets, the outcomes were even more variable, with the geminate clade clustering together in 71% and 86% of random draws using the BCD and unweighted UniFrac metrics, respectively. This highlights two important points. First, although rarefaction is commonly used, and often recommended [28,45], for standardization of microbiome data, our results show that in some cases, hypothesis testing using rarefied datasets can produce different results than other normalization approaches. Second, using a single rarefied ASV table to test hypotheses, as is common in microbial ecology, can be problematic given the stochastic nature of individual rarefaction draws. Instead, we recommend using multiple iterations to test the robustness of results using rarefied data. Finally, our network analyses identified multiple taxa that helped differentiate the geminate clade from the outgroup, *C. montagnei*, but none of these showed significant co-phylogenetic relationships with the host group. Taken together, our results suggest that in the *Cerithideopsis* whole-body samples, host filtering and/or host ecology likely drive any observed similarity between the microbiomes of the geminate species. As stated above, the climbing behavior of *C. montagnei* may limit its exposure to seawater microbes thus differentiating the microbiome composition of this species from its entirely mud-dwelling relatives.

In contrast to the whole-body samples, shell-surface microbiomes of *Cerithideopsis* species consistently mirrored the host phylogeny. Several microbial clades were responsible for this differentiation between the geminate species and *C. montagnei*, some of which showed co-phylogenetic relationships with these hosts. These include members of the microbial orders Thermoanaerobaculales and SBR1031, and the class Gammaproteobacteria. Thermoanaerobaculales is an order within the phylum Acidobacteriota, recently described from freshwater hot springs [46]. Little is currently known about this clade, but members have been found in high abundance in intertidal sands [47] and are present in many of our seawater and sediment samples, suggesting that these microbes may be transferred from the environment to shell-surfaces. SBR1031 is an order of nitrifying bacteria within the phylum Chloroflexi previously found in association with marine invertebrates [48,49] and the mangrove rhizosphere [50]. Since shell-surfaces are sites of active nitrogen cycling and nitrification in the marine environment [38,39], and members of this order were present in our sediment and seawater samples, this association may also constitute an acquisition from the environment. Gammaproteobacteria is a highly diverse clade common in the marine environment and many marine hosts [40,51], so in this case the processes driving a co-phylogenetic pattern with *Cerithideopsis* are unclear. In general, it is surprising that significant co-phylogenetic divergence between geminate hosts and their microbiomes was only evident in shell-surface biofilms of *Cerithideopsis*, as these communities are expected to be more influenced by the external environment than the biology of host [16,52].

While the formation of the IP led to predictable patterns of evolutionary divergence of marine animal species on either side, our results show that this geologic event affected host-associated microbiomes in a much more complex and context-dependent manner. Depending on the geminate clade and host tissue sampled, we found evidence for (i) phylosymbiosis driven by environmental filtering, (ii) co-phylogenetic divergence of hosts and their microbial associates and (iii) no discernable impact of host evolutionary history on microbiomes of geminate species. These results could reflect multiple causes. First, an expectation of co-divergence between hosts and their microbiomes due to the rise of the IP assumes that populations of individual microbial taxa were originally present in both oceans and subsequently split by the closure of the IP, leaving representatives on both sides. However, microbiome compositions of marine invertebrate hosts commonly differ substantially between populations of the same host species [e.g., 36,46] as shown by our samples of *Cerithium stercusmuscarum*. If populations of hosts did not share many microbes prior to being separated by the IP, it is perhaps unsurprising that observed phylosymbiosis patterns are driven by a small number of taxa, rather than being a community-wide trend. Second, for co-phylogenetic divergence of hosts and their microbes to be detectable millions of years after a vicariant event, hosts and microbial taxa must diverge at similar rates. However, due to their short generation times and differences in selective pressures between the Caribbean and Eastern Pacific after the closure of the IP, the divergence rates of microbial taxa in separate oceans were likely much higher than those of their hosts. Such differences in evolutionary rates can decrease the phylosymbiosis signal and create closer evolutionary relationships between microbial taxa currently present on the same side of the IP, as we found in multiple cases. Unfortunately, current data limitations make this difficult to test, as ancestral distributions of microbial taxa remain unknown and calibrating microbial phylogenies also remains challenging [53].

Overall, our results suggest that ecological and evolutionary responses of host-associated microbial communities to major vicariant events such as the rise of the Isthmus of Panama, are complex and taxon-specific. Future comparative analyses of microbiomes of other geminate hosts would be useful for better understanding how the microbiome compositions of marine hosts have been shaped by major geological events.

## Acknowledgements

We thank E. Caballero and G. Castellanos-Galindo for assistance with sampling and C. Schloeder, I. Ochoa and R. Collin, for sample storage, shipping and logistical support. ATN was funded by a STRI Short-Term Research Fellowship. All samples were collected in compliance with permits #SC/A-1-19 (collections) and #SEX/A-33-19 (export) from the Ministerio de Ambiente de Panamá.

## Data Accessibility Statement

16S rRNA gene sequence data and associated metadata are available from the National Center for Biotechnology Information (NCBI) Sequence Read Archive (SRA) under Bioproject accession number PRJNA744215. The ASV tables used for analyses will be made available via Dryad.

