## Supplemental Tables and Figures for "Microbiome divergence of marine gastropod species separated by the Isthmus of Panama"

Supplement 1

**Table of Contents:**

| **Table S1** | Page 2 |
| --- | --- |
| **Table S2** | Page 2 |
| **Table S3** | Page 3 |
| **Table S4** | Page 3 |
| **Table S5** | Page 4 |
| **Table S6** | Page 4 |
| **Figure S1** | Page 5 |
| **Figure S2** | Page 6 |
| **Figure S3** | Page 7 |
| **Figure S4** | Page 8 |
| **Figure S5** | Page 9 |
| **Figure S6** | Page 9 |
| **Figure S7** | Page 10 |

Table S1. The number of whole-body and shell-surface samples collected from each host species at each site.

| Host species | Collection site | Whole-body samples (*n*) | Shell-surface samples (*n*) |
| --- | --- | --- | --- |
| *Cerithium atratum* | Punta Galeta | 5 | 7 |
| *Cerithium lutosum* | Punta Galeta | 6 | 7 |
| *Cerithium stercusmuscarum* | Bique | 7 | 6 |
| *Cerithium stercusmuscarum* | Punta Culebra | 5 | 5 |
| *Cerithideopsis mazatlanica* | Bique | 8 | 6 |
| *Cerithideopsis montagnei* | Bique | 6 | 6 |
| *Cerithideopsis pliculosa* | Puerto Pilón | 8 | 6 |
| *Cerithideopsis valida* | Bique | 7 | 5 |

Table S2. Results of Kruskal-Wallis tests comparing Hill numbers at q = 0 and q = 1 across species within each host genus and body site, based on the unrarefied datasets.

| Sample Type | Diversity Metric | X^2^ | p |
| --- | --- | --- | --- |
| *Cerithium* whole-body | q = 0 | 1.200 | 0.273 |
| *Cerithium* whole-body | q = 1 | 11.858 | **0.0079** |
| *Cerithium* shell-surface | q = 0 | 0.0041 | 0.949 |
| *Cerithium* shell-surface | q = 1 | 2.159 | 0.142 |
| *Cerithideopsis* whole-body | q = 0 | 9.902 | **0.0194** |
| *Cerithideopsis* whole-body | q = 1 | 6.203 | 0.102 |
| *Cerithideopsis* shell-surface | q = 0 | 6.541 | 0.088 |
| *Cerithideopsis* shell-surface | q = 1 | 6.451 | **0.0048** |

Table S3. Results of Kruskal-Wallis tests comparing observed ASV richness and Shannon’s H’ across species within each host genus and body site, based on the average of 100 datasets rarefied to 3,000 sequences/sample.

| Sample Type | Diversity Metric | X^2^ | p |
| --- | --- | --- | --- |
| *Cerithium* whole-body | Observed ASVs | 5.483 | 0.251 |
| *Cerithium* whole-body | Shannon’s H’ | 6.818 | **0.009** |
| *Cerithium* shell-surface | Observed ASVs | 0.033 | 0.855 |
| *Cerithium* shell-surface | Shannon’s H’ | 1.633 | 0.201 |
| *Cerithideopsis* whole-body | Observed ASVs | 7.531 | 0.057 |
| *Cerithideopsis* whole-body | Shannon’s H’ | 6.436 | 0.092 |
| *Cerithideopsis* shell-surface | Observed ASVs | 7.662 | 0.053 |
| *Cerithideopsis* shell-surface | Shannon’s H’ | 9.517 | **0.023** |

Table S4. PERMANOVA and betadisper results from all sample types, normalized by total sum scaling, using both Bray-Curtis dissimilarity and unweighted UniFrac distance. A significant betadisper p value in the *Cerithium* whole-body samples was found due to the divergence between the two populations of *C. stercusmuscarum*; though after removing one population from analysis, species were still found to be significantly different from one another (PERMANOVAs: all *p < 0.001*, betadisper: all *p > 0.2*). In the *Cerithideopsis* whole-body samples, a significant betadisper p value is unlikely to impact the interpretation of the PERMANOVA results, as PERMANOVA is robust to heterogeneity of dispersions when sample sizes are similar.

| Sample Type | Diversity Metric | F Model | R^2^ | p | betadisper |
| --- | --- | --- | --- | --- | --- |
| *Cerithium whole-body* | Bray-Curtis | 5.226 | 0.343 | **<0.001** | 0.969 |
| *Cerithium whole-body* | Unweighted UniFrac | 1.802 | 0.153 | **<0.001** | **0.005** |
| *Cerithium shell-surface* | Bray-Curtis | 3.537 | 0.243 | **<0.001** | 0.510 |
| *Cerithium shell-surface* | Unweighted UniFrac | 1.660 | 0.131 | **<0.001** | 0.902 |
| *Cerithideopsis whole-body* | Bray-Curtis | 3.484 | 0.295 | **<0.001** | **<0.001** |
| *Cerithideopsis whole-body* | Unweighted UniFrac | 2.082 | 0.200 | **<0.001** | 0.086 |
| *Cerithideopsis shell-surface* | Bray-Curtis | 4.149 | 0.396 | **<0.001** | 0.466 |
| *Cerithideopsis shell-surface* | Unweighted UniFrac | 1.475 | 0.189 | **<0.001** | 0.455 |

Table S5. PERMANOVA and betadisper results from all sample types, normalized by rarefying to 3,000 sequences per sample, using both Bray-Curtis dissimilarity and unweighted UniFrac distance.

| Sample Type | Diversity Metric | F Model | R^2^ | p | betadisper |
| --- | --- | --- | --- | --- | --- |
| *Cerithium* whole-body | Bray-Curtis | 5.392 | 0.362 | **<0.001** | 0.278 |
| *Cerithium* whole-body | Unweighted UniFrac | 1.946 | 0.170 | **<0.001** | **<0.001** |
| *Cerithium* shell-surface | Bray-Curtis | 4.345 | 0.367 | **<0.001** | 0.097 |
| *Cerithium* shell-surface | Unweighted UniFrac | 2.151 | 0.223 | **<0.001** | 0.218 |
| *Cerithideopsis* whole-body | Bray-Curtis | 3.191 | 0.277 | **<0.001** | **<0.001** |
| *Cerithideopsis* whole-body | Unweighted UniFrac | 2.204 | 0.209 | **<0.001** | 0.054 |
| *Cerithideopsis* shell-surface | Bray-Curtis | 3.971 | 0.385 | **<0.001** | 0.397 |
| *Cerithideopsis* shell-surface | Unweighted UniFrac | 1.675 | 0.209 | **<0.001** | 0.714 |

Table S6. PERMANOVA and betadisper results comparing host species to environmental samples, normalized by total sum scaling, using both Bray-Curtis dissimilarity and unweighted UniFrac distance.

| Sample Type | Diversity Metric | F Model | R^2^ | p | betadisper |
| --- | --- | --- | --- | --- | --- |
| *Cerithium* whole-body | Bray-Curtis | 5.483 | 0.174 | **<0.001** | 0.283 |
| *Cerithium* whole-body | Unweighted UniFrac | 3.026 | 0.074 | **<0.001** | 0.988 |
| *Cerithium* shell-surface | Bray-Curtis | 5.297 | 0.117 | **<0.001** | 0.288 |
| *Cerithium* shell-surface | Unweighted UniFrac | 3.014 | 0.070 | **<0.001** | 0.205 |
| *Cerithideopsis* whole-body | Bray-Curtis | 6.315 | 0.126 | **<0.001** | 0.536 |
| *Cerithideopsis* whole-body | Unweighted UniFrac | 2.667 | 0.057 | **<0.001** | 0.725 |
| *Cerithideopsis* shell-surface | Bray-Curtis | 5.283 | 0.122 | **<0.001** | 0.417 |
| *Cerithideopsis* shell-surface | Unweighted UniFrac | 2.133 | 0.053 | **<0.001** | 0.080 |

Table S7. Microbial taxa at each taxonomic level found to be significantly associated with a geminate species group by SPLS-DA analysis. Bolded taxa are those presenting potential cophylogenetic patterns with the host clade using parafit (*p < 0.05*). An “X” indicates that no significant associations were found.

| Sample type | ASV | Genus | Family | Order | Class |
| --- | --- | --- | --- | --- | --- |
| *Cerithium* whole-body | ASV10677 | X | X | X | X |
| *Cerithium* shell-surface | X | X | X | X | X |
| *Cerithideopsis* whole-body | X | *Roseibacillus*  *Roseibacterium* | DEV007  Gimesiaceae  Pirellulaceae  Rubinisphaeraceae | X | X |
| *Cerithideopsis* shell-surface | X | *Actibacterium*  *Aquimarina*  *Granulosicoccus*  *Sva0081* | **A4b**  DEV007  Microtrichaceae  **Thermoanaerobaculaceae**  Xenococcaceae | Ardenticatenales  **SBR1031**  Steroidobacterales  **Thermoanaerobaculales** | Desulfobacteria  **Gammaproteobacteria** |

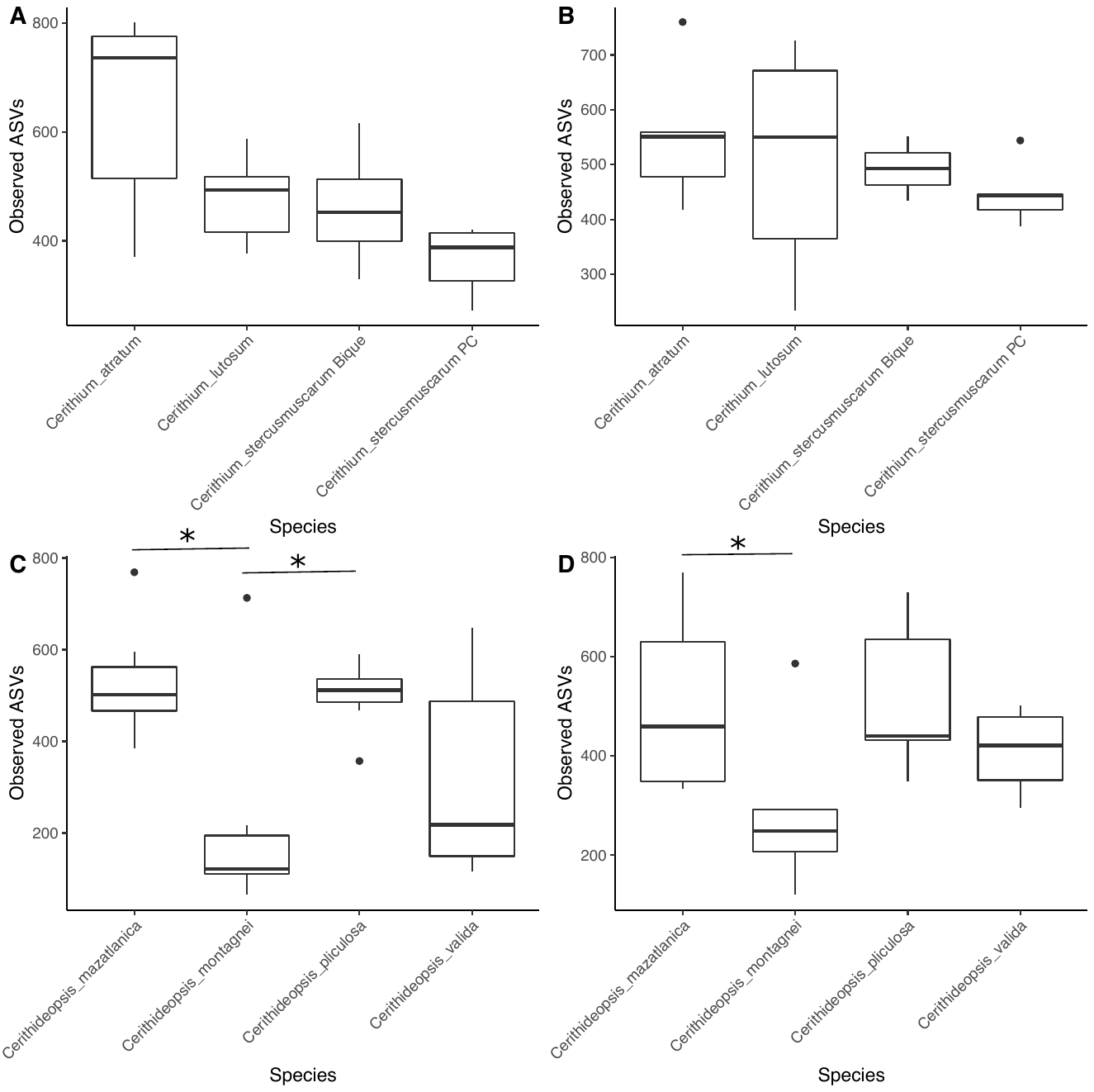

Figure S1. Observed ASV richness, based on the average of 100 ASV tables rarefied to 3,000 sequences per sample, for *Cerithium* (A,B) and *Cerithideopsis* (C,D) whole-body (A,C) and shell-surface (B,D) samples. Asterisks indicate samples which were found to be significantly different by Dunn’s test at p < 0.05 (*).

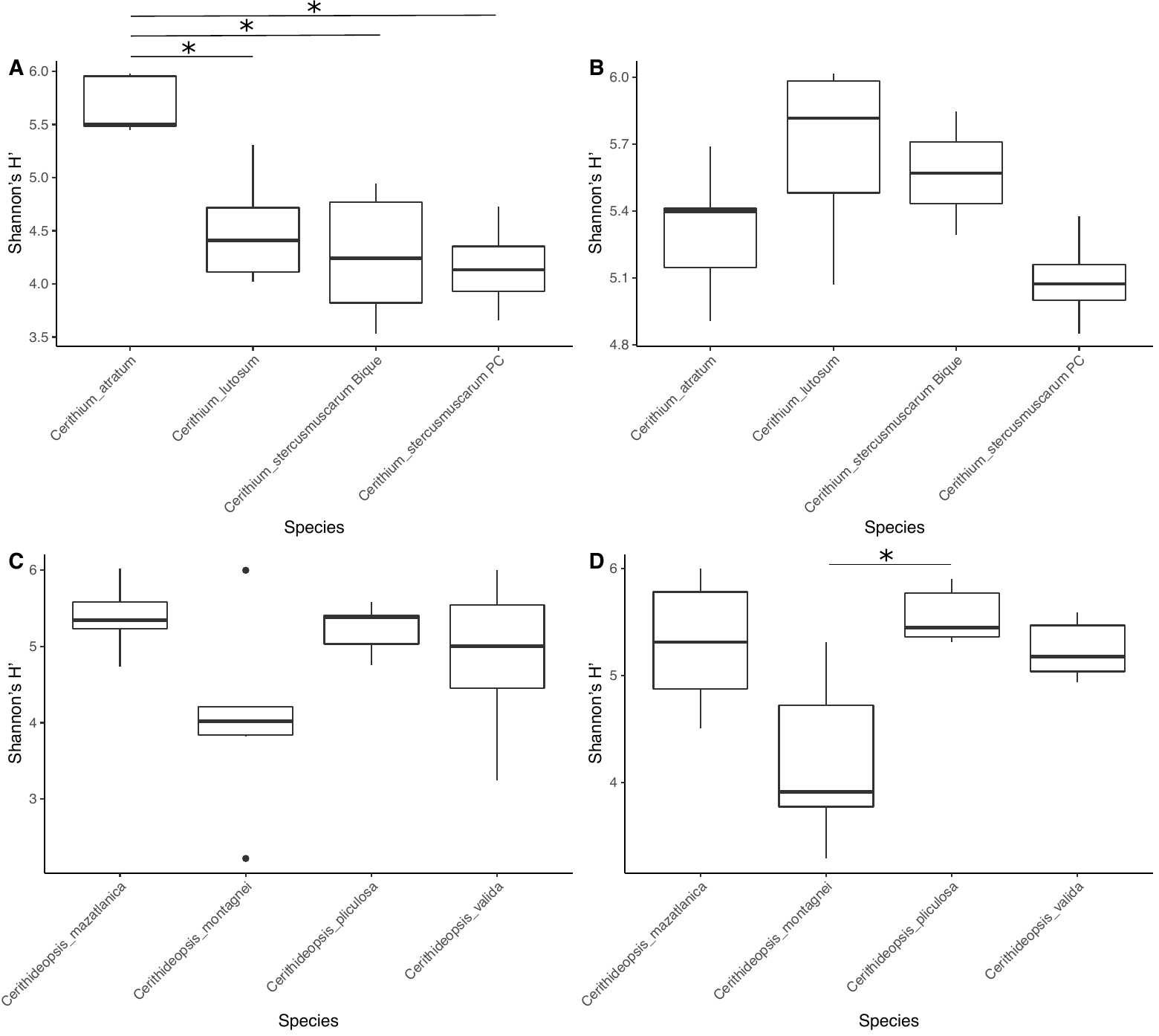

Figure S2. Shannon’s H, based on the average of 100 ASV tables rarefied to 3,000 sequences per sample, for *Cerithium* (A,B) and *Cerithideopsis* (C,D) whole-body (A,C) and shell-surface (B,D) samples. Asterisks indicate samples which were found to be significantly different by Dunn’s test at p < 0.05 (*).

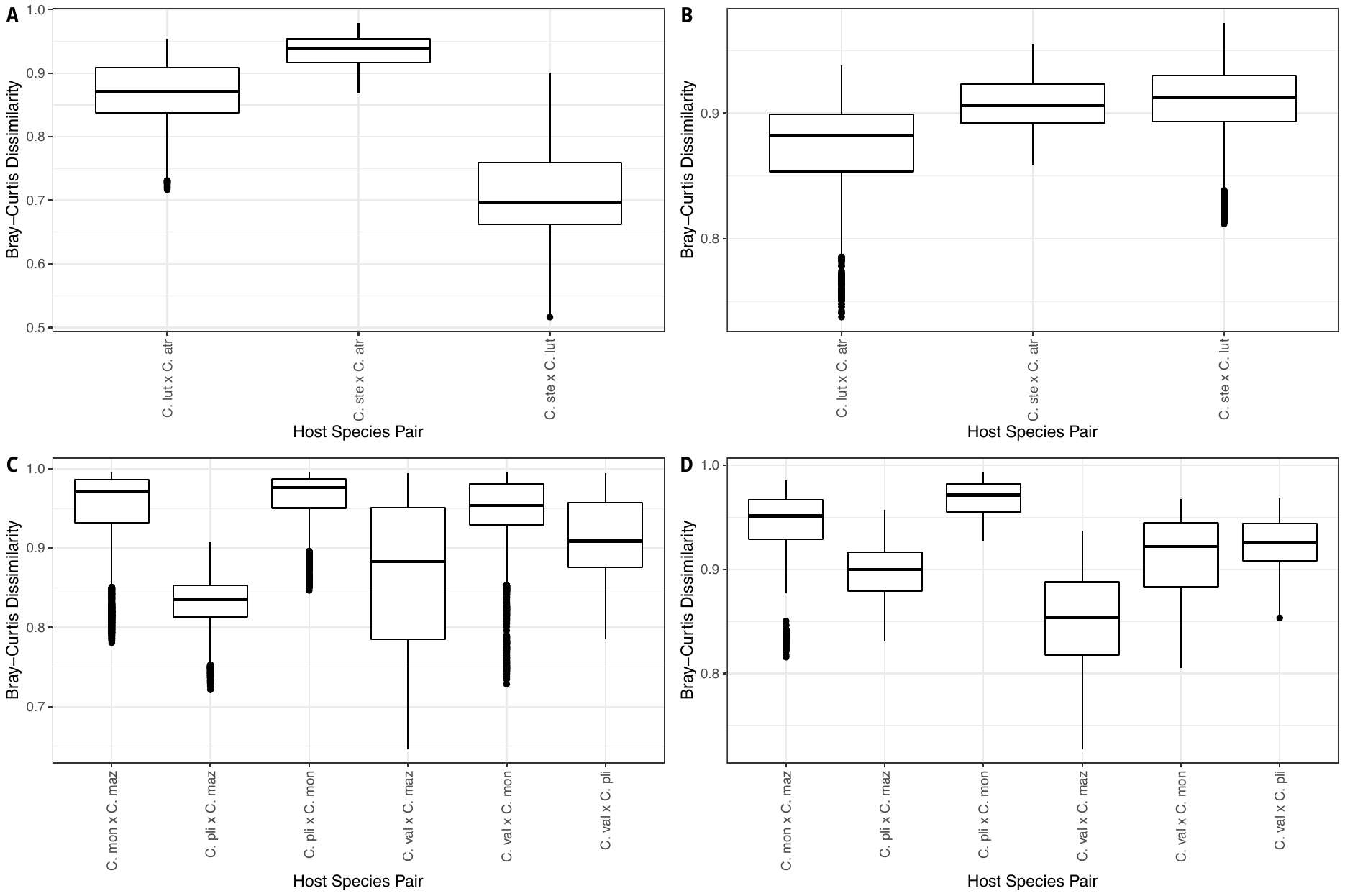

Figure S3. Boxplots of Bray-Curtis dissimilarity between each host species pair in the *Cerithium* (A,B) and *Cerithideopsis* (C,D) whole-body (A,C) and shell-surface (B,D). Boxplots show all values across the 100 datasets rarefied to 3,000 sequences/sample.

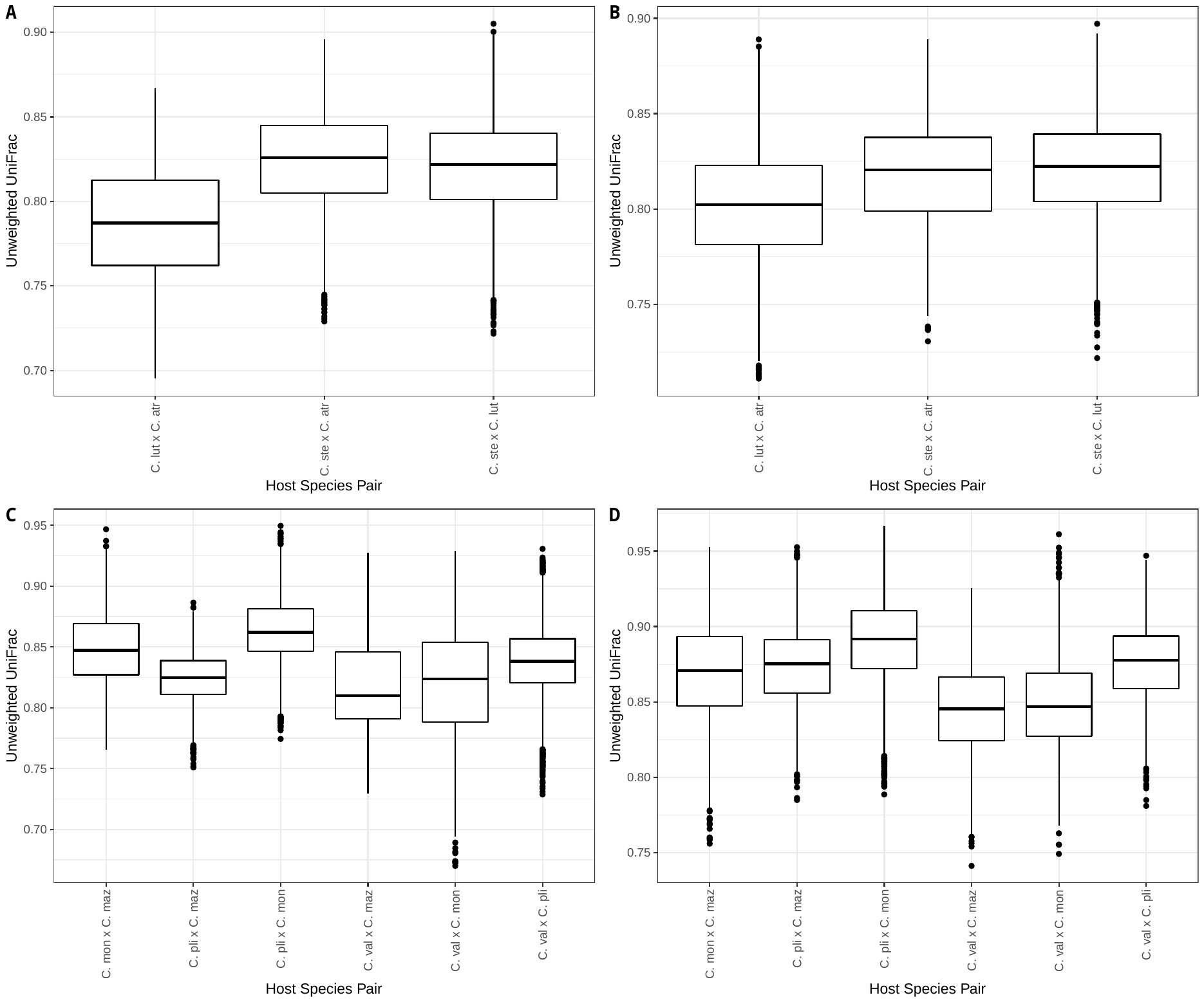

Figure S4. Boxplots of unweighted UniFrac distance between each host species pair in the *Cerithium* (A,B) and *Cerithideopsis* (C,D) whole-body (A,C) and shell-surface (B,D). Boxplots show all values across the 100 datasets rarefied to 3,000 sequences/sample.

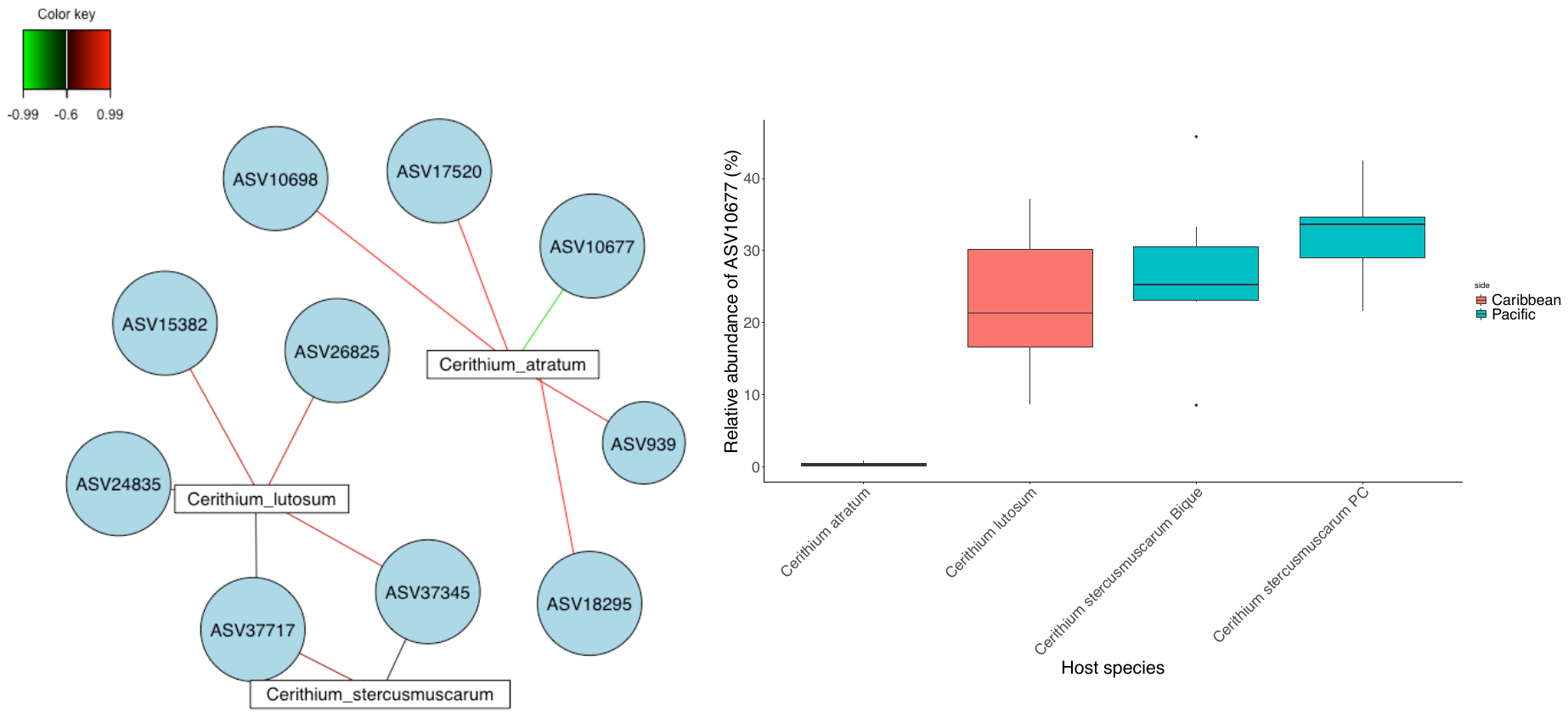

Figure S5. SPLS-DA network of ASVs (left) and relative abundance of ASV10677 (right) in *Cerithium* whole-body samples.

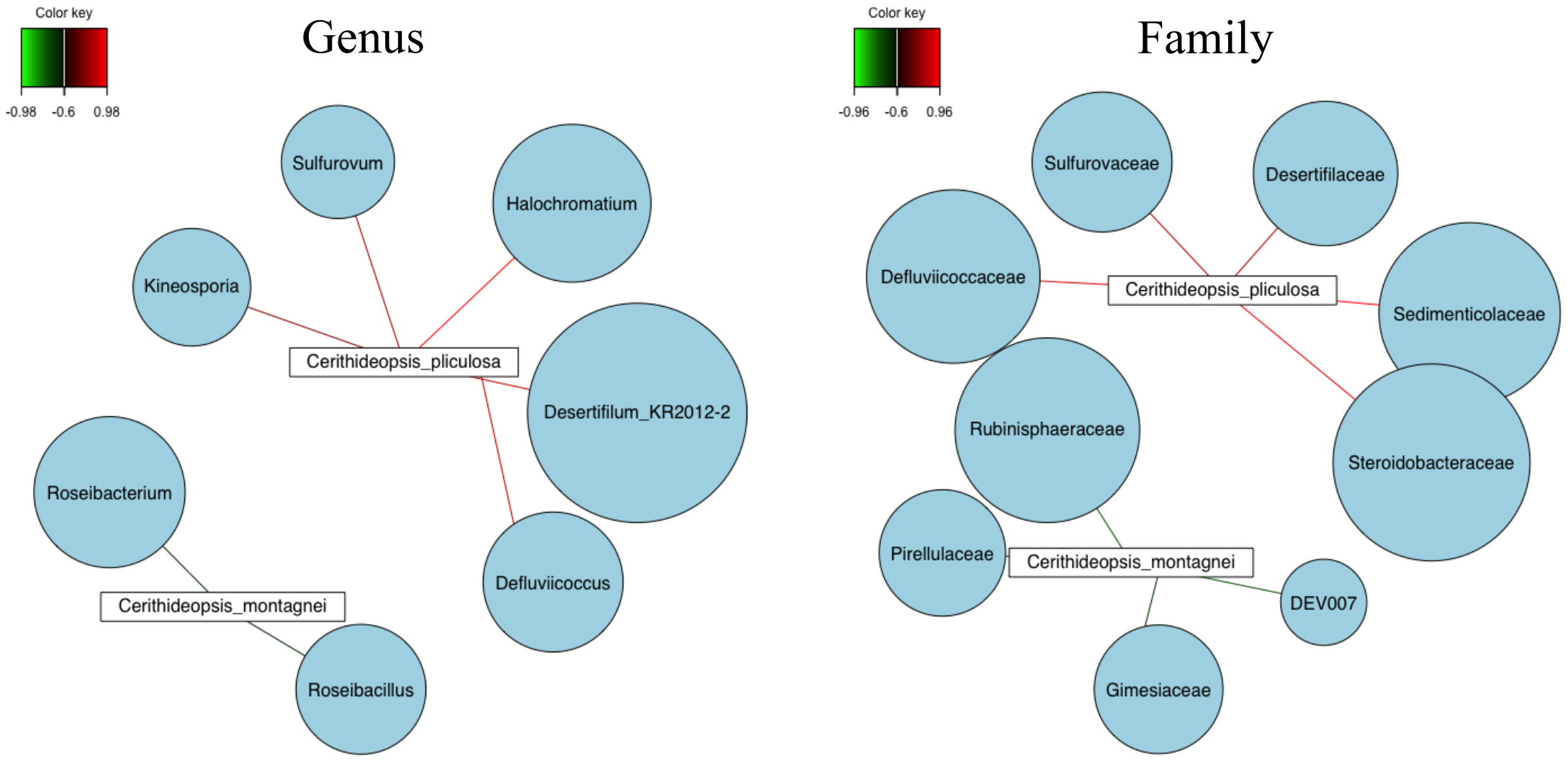

Figure S6. SPLS-DA networks of microbial genera (left) and families (right) in *Cerithideopsis* whole-body samples.

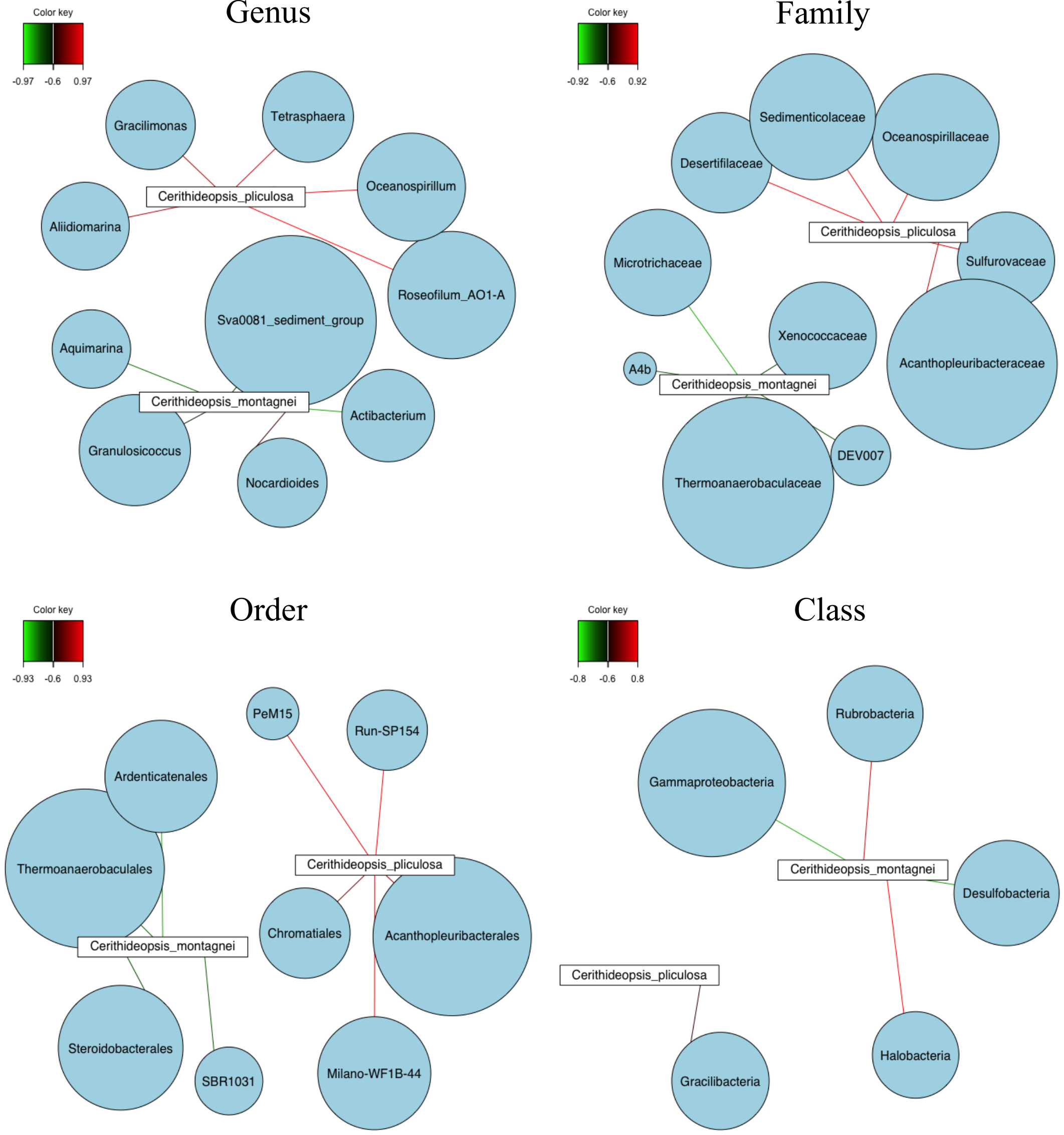

Figure S7. SPLS-DA networks of microbial genera (top left), families (top right), orders (bottom left) and classes (bottom right) in *Cerithideopsis* shell-surface samples.
